# Stiffness-tunable neurotentacles for minimally invasive implantation and long-term neural activity recordings

**DOI:** 10.1101/2024.07.21.604464

**Authors:** Yang Wang, Xing Xu, Xiaowei Yang, Rongyu Tang, Ying Chen, Yijun Wang, Jing Liang, Weihua Pei

**Author notes:** Corresponding author. Email: Weihua Pei or Jing Liang. These authors contributed equally to this work: Yang Wang, Xing Xu.

## Abstract

Flexible implantable microelectrodes have been demonstrated to exhibit excellent biocompatibility for chronic neural activity recordings. However, the low bending strength of the commonly employed flexible materials presents a significant challenge for probe insertion into the brain. Traditional implantation methods for flexible electrodes generally require additional auxiliary materials or tools, which tend to have a much larger footprint than the probes themselves, greatly increasing the damage to neurons during insertion. Here we have proposed a stiffness-tunable polyimide probe for deep brain implantation, referred to as Neurotentacle, enabled by embedded microchannels in which the liquid pressure is controllable (from 0.1MPa to more than 2.0MPa). During the insertion phase into the brain, the neurotentacle can pose a high stiffness under elevated internal pressure to penetrate the brain tissues without the use of any additional materials or tools. Once the device has been successfully inserted, it can regain its flexibility by reducing the internal pressure. Importantly, the novel multilayer microfabrication process keeps the structural dimensions of the neurotentacle similar to those of a regular flexible probe. Therefore, the neurotentacle can produce an extremely low level of damage to brain tissue during its insertion phase, while extending its long-term biocompatibility and stability, which has been experimentally verified in histological evaluations conducted on both acute and chronic animal specimens. In addition, the chronically implanted neurotentacles enabled stable neural activity recordings in mice with an average spike yield of 96% and an average signal-to-noise ratio of 15.2. The proposed neurotentacle does not necessitate the use of complex devices and its insertion process is straightforward and highly controllable, thus rendering it an appealing technique for minimally invasive implantation and long-term neural recording of flexible electrodes.

## 1. Introduction

Implantable microelectrodes are capable of acquiring electrophysiological signals of neurons with high temporal-spatial resolution, which is of great importance in brain-computer interface research[1]. In particular, the flexible neural microelectrodes have been demonstrated to exhibit excellent biocompatibility, which effectively reduces the immune response of brain tissue and extends the in vivo longevity of the device[2–4]. A variety of flexible electrodes have been developed based on different materials, such as Polyimide (PI)[5–7], Parylene[8, 9], SU-8[10, 11], and so on. However, the low Young’s modulus of these materials implies low bending stiffness for most of the flexible probes[12, 13], which presents a challenge for their implantation into the brain.

A variety of implantation methods have been developed to facilitate the delivery of flexible electrodes into the brain. These methods can be broadly divided into two categories: methods of reinforcing flexible electrodes with sacrificial materials (RFSM methods) and methods of inserting flexible electrodes with auxiliary tools (IFAT methods)[14]. The RFSM methods typically strengthen the flexible probes by employing biodegradable materials, including gelatin[15], maltose[16], dextran[17], polyethylene glycol (PEG)[3, 18], polyacetate[19], silk proteins[20], and some other polymers with similar properties[21]. These materials can increase the stiffness of the probes prior to implantation, enabling them to penetrate the brain. Following implantation, the degradation of these materials within the brain allows the probes to recover their flexibility. However, the footprint of the reinforced flexible electrode is usually much larger than the original, enlarging the acute damage caused by the probe insertion. In addition, these materials dissolve rapidly under physiological conditions, making it difficult to manipulate the insertion speed. The IFAT methods temporarily attach the flexible probes to a rigid tool such as a silicon shuttle or metal microneedle and then withdraw the tool after the electrodes are inserted. The attachment can be achieved by self-adsorption effect[22], water-soluble adhesive bonding[23], needle-hole assembling[11, 24, 25], syringe holding[26], and so on. However, these auxiliary tools are also much larger than the flexible electrodes themselves. Moreover, they could cause secondary damage to brain tissue when withdrawn, as well as disturb or even bring out the inserted flexible electrodes.

In addition, to minimize insertion damage, the optimal implantation should not involve the use of any auxiliary tools or materials that would enlarge the flexible electrodes to an excessive degree. Some novel insertion methods, such as magnetic actuation[27] and microfluidic actuation[28], have the potential to meet this need. Nevertheless, these methods still require optimization in terms of drive force, device complexity, insertion speed, and other aspects. Some strength-adaptive materials can be integrated with the electrode substrate, such as nanocomposites[29], shape memory polymers[30], and liquid crystal polymers[31]. These materials are sufficiently stiff to penetrate brain tissue before insertion and will gradually soften under physiological conditions after insertion. However, such materials typically require complex processing procedures to avoid the conditions that could denature them, such as the high temperatures or some chemical solution environments common in MEMS. Moreover, the bending stiffness of these materials is still insufficient to support very small and thin probes.

In this study, we designed and prepared a neurotentacle probe with tunable stiffness based on liquid pressure. No auxiliary materials or tools other than saline were employed for the insertion of the neurotentacle into the brain. Inside the neurotentacle, there is a microchannel, through which the liquid can be injected and the pressure can be regulated to adjust the stiffness of the probe. Based on the developed ultra-thin microchannel process, the encapsulated neurotentacles had comparable footprints to the regular flexible probes, as evidenced by previous literature[6, 24, 25, 32, 33]. There was only a slight expansion of the neurotentacle volume during insertion, and no further damage was caused by retracting the auxiliary tools, thus greatly reducing the acute damage. In addition, chronically implanted neurotentacles enabled long-term stable neural activity recordings in mice with a high yield of spikes and high signal-to-noise ratios (SNR).

## 2. Results

### 2.1 Design of neurotentacles

The proposed neurotentacle probe can change its strength as the internal pressure varies. This allows it to increase the strength to penetrate brain tissue before implantation and decrease the strength to recover flexibility after implantation, as shown in **Fig.1A** and **Movie S1**. The key to neurotentacle design is the integration of a microchannel into the typical PI flexible probe. Compared with the common sandwich-type PI probes (PI-Metal-PI)[6, 24], the neurotentacle has an additional microchannel in the middle of the insulating layer (PI-Microchannel-PI-Au-PI), as shown in **Fig.1B**. At the rear end of the neurotentacle, there reverses an inlet for the liquid to enter the microchannel. A liquid injection device is connected to the inlet via a pipeline. It can pump liquid into the probe and regulate its internal pressure. A hydrometer is mounted on the injection device for monitoring the liquid pressure in real-time.

**Fig 1.**
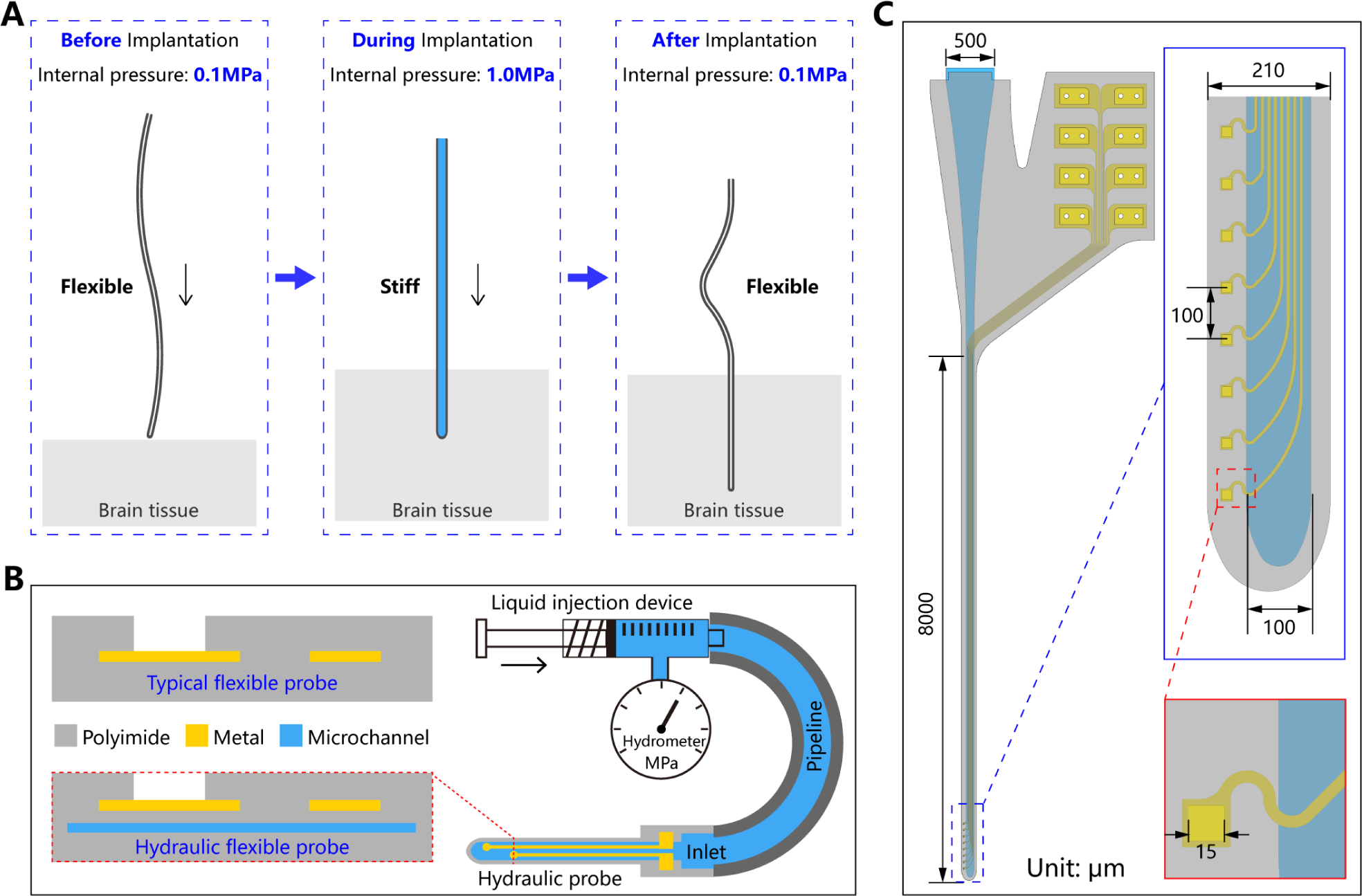
Design of neurotentacles. (A) Implantation process of the neurotentacle based on variable stiffness. (B) Schematic diagram of the overall design and packaging of the neurotentacle. (C) Layout design of the neurotentacle.

The layout of the neurotentacle is shown in **Fig.1C**. The shank is 8 mm long, enough to cover the entire brain area of small rodents. It has a width of 210 μm, of which the microchannel width is 100 μm. At the front end of the probe, there are eight recording sites with a diameter of 15 μm and a center-to-center distance of 80 μm. The metal traces are above the microchannel, and all the recording sites are arranged at the edge of the shank. This layout is partly because it is easier to capture neuronal signals at the edges, and partly to avoid over-etching to break the microchannel. The metal traces are designed to be S-shaped at the edge of the microchannel to enhance their tensile resistance, thus better adapting to the variable morphology of the probe during the filling and emptying of the liquid. At the back end of the neurotentacle, the liquid inlet and the metal pads are separated to avoid subsequent conflicts between the operations of pipeline encapsulation and electrical connection.

### 2.2 Fabrication of neurotentacles

The preparation of the microchannel is the most important part of fabricating the neurotentacle. It is related to many aspects such as process complexity, microchannel intensity, probe thickness, and encapsulation difficulty. Here, we proposed a novel process flow for fabricating the ultra-thin microchannel based on differences in adhesion, as shown in **Fig.2A**. Reactive ion etching and hydrophobic treatment were employed to create regions with different adhesion on the first PI layer. When the second PI layer was cured, the strongly adhesive regions bonded together and the weakly adhesive regions formed the microchannel precursor. Once the neurotentacle is encapsulated, the liquid injected into the microchannel precursor can open it up, thereby forming the microchannel. The microchannel prepared based on this method is characterized by an extremely small volume in the unfilled state, which greatly reduces the overall thickness of the probe. Its preparation process is easy to realize and involves only the materials and equipment commonly used to fabricate PI-based flexible probes.

**Fig 2.**
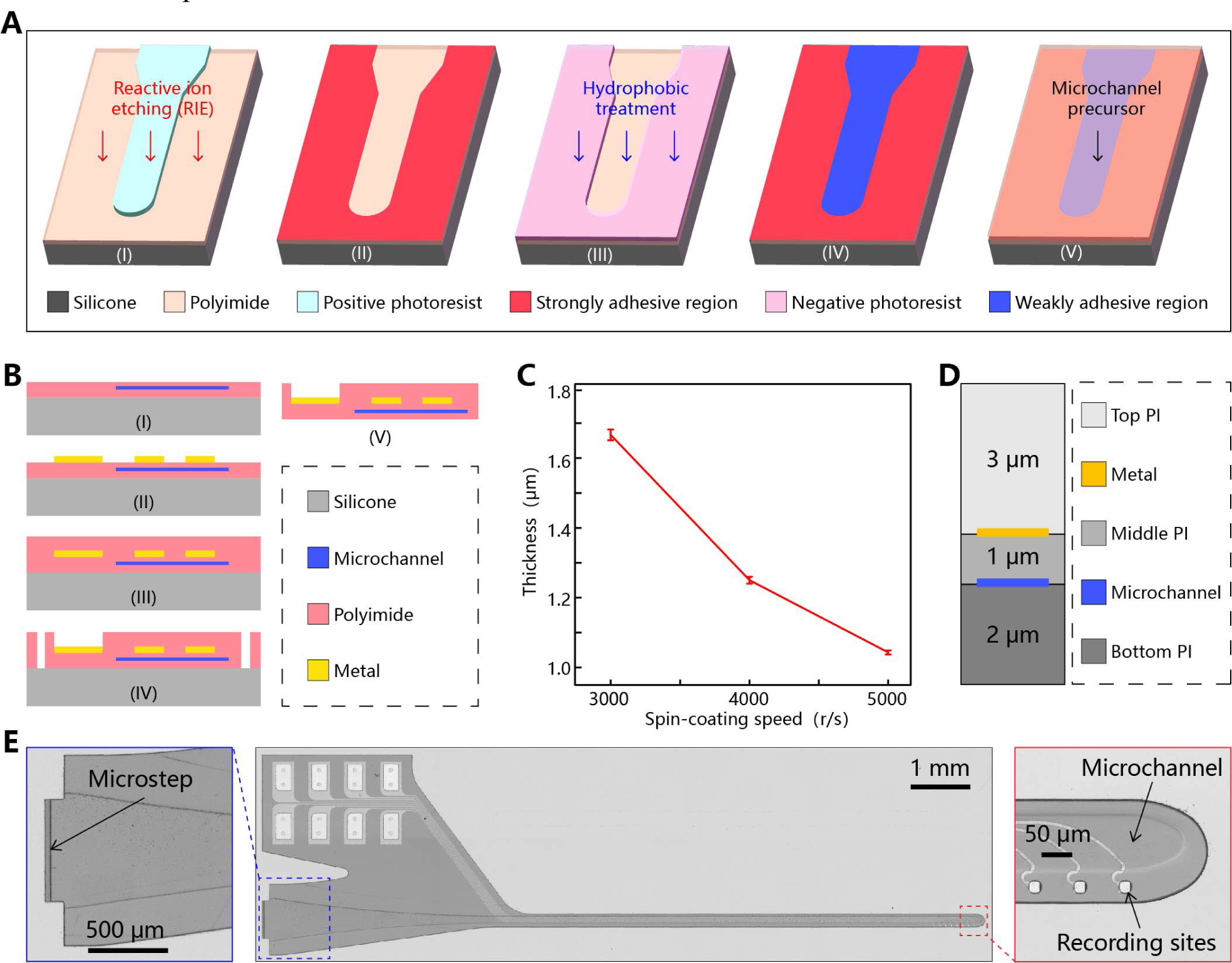
Fabrication of neurotentacles. (A) Process flow for the preparation of ultra-thin microchannel. (B) Procedures for fabricating the flexible probe after constructing the microchannel. (C) The relationship between spin-coating speed and film thickness for diluted PI2611. (D) Optimized thickness distribution of three PI layers of the neurotentacle. (E) A prepared neurotentacle probe based on the MEMS process (Middle). Liquid inlet at the end of the probe (Left). The tip of the probe with visible microchannel and recording sites(Right).

After forming the microchannel precursor, the neurotentacle is fabricated in the same process as a conventional PI probe, as shown in **Fig.2B**. The presence of the microchannel renders the device to contain three layers of PI. The method of thinning the PI membrane[34] is employed herein to keep the overall thickness of the neurotentacle at the same level as that of regular PI electrodes. PI2611 was mixed with NMP in a mass ratio of 4:1 to obtain the diluted PI. By controlling the spin-coating speed, the PI films can be prepared with different thicknesses from 1 to 2 μm. The relationship between the thickness and the spin-coating speed is shown in **Fig. 2C**. It should be noted that the probe thickness is not equally proportional for each layer of PI. The thickness distribution of the top, middle, and bottom PI layers needs to be optimized for two reasons. Firstly, it is necessary to ensure that the PI thicknesses above and below the metal layer are uniform to avoid self-bending of the released probes (**Fig.S1**). This means that the total thickness of the middle and bottom PIs should ideally be the same as that of the top PI. Secondly, the bottom PI should be as thick as possible to avoid breaking the microchannel during encapsulation. The optimized PI thickness is shown in **Fig. 2D**. The prepared neurotentacle is shown in **Fig. 2E**. The total thickness was about 6.2 μm, with the PI above and below the metal layer being 3.3 and 2.9 μm, respectively. Such a neurotentacle is as thin as the typical PI-based flexible probe[6, 25].

### 2.3 Encapsulation of neurotentacles

To facilitate the opening of the microchannel, a micro-step is reserved at the end of the probe (**Fig.2E**). Along the bottom of the micro-step, the PI can be split apart by using a sharp object, thus forming the inlet of the microchannel (**Fig.3A**). A stainless tube was inserted into the microchannel as the relay. Then a silicone tube and a syringe head were sequentially connected to the microchannel, as shown in **Fig.3B**. All the joints were sealed with ultraviolet (UV) glue. During the encapsulation of the liquid pathway, the neurotentacle was also electrically encapsulated with the printed circuit board (PCB) and connector. The detailed encapsulation procedures (**Fig.S2**) can be found in Materials and Methods.

**Fig 3.**
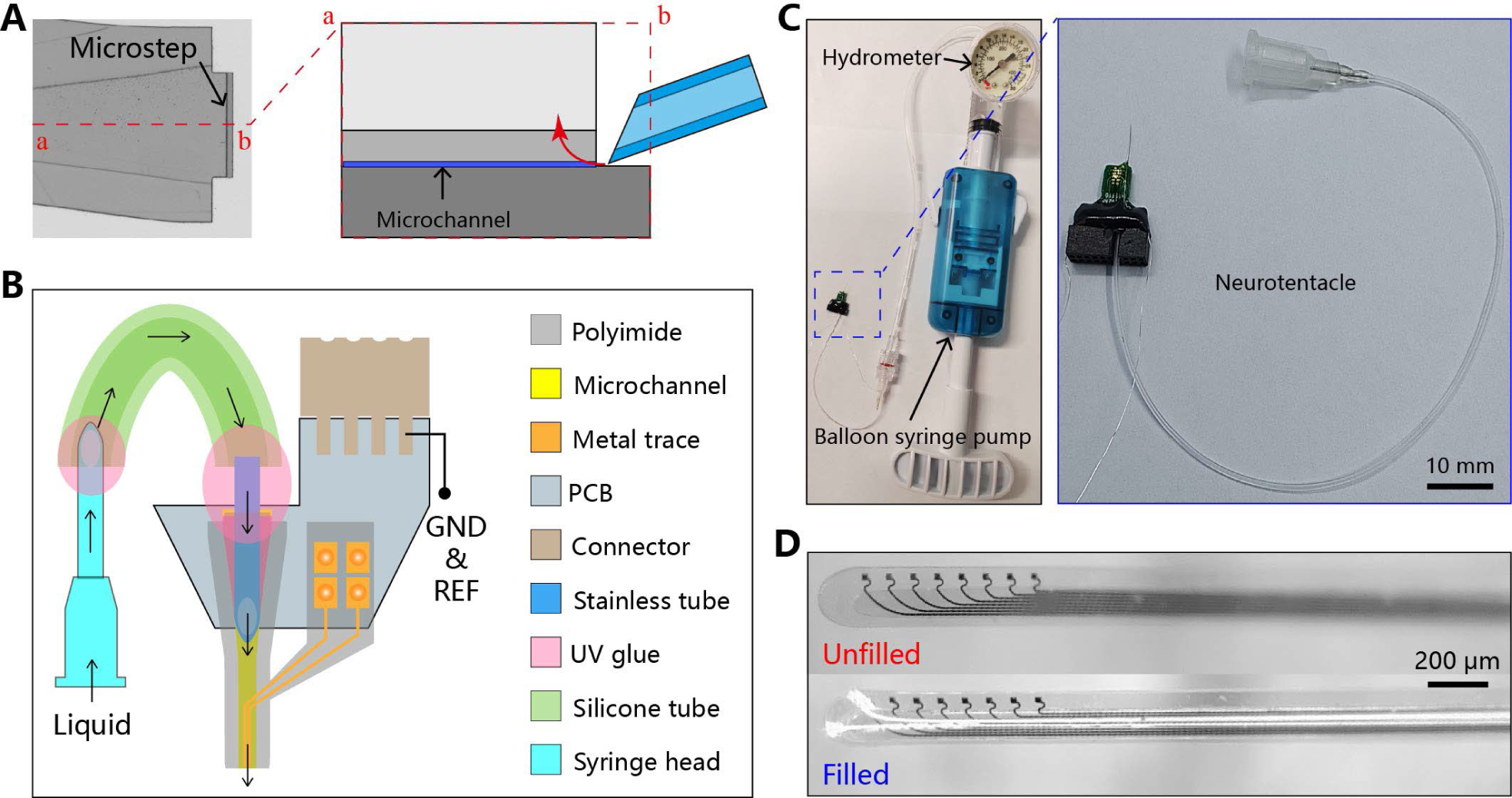
Encapsulation of neurotentacles. (A) Schematic diagram of opening the liquid inlet through the micro-step. (B) Schematic diagram of encapsulating the neurotentacle with the liquid pathway and electrical connector. (C) A physically encapsulated neurotentacle and a syringe pump used to apply the hydraulic pressure. (D) Comparison of the morphology of a neurotentacle under the filled and unfilled state.

A fully encapsulated neurotentacle is shown in **Fig.3C**. A medical Balloon Syringe Pump was used to inject liquid into the neurotentacle. It can provide a stable pressure for the neurotentacle, and the hydrometer on it can monitor the pressure in real-time with a maximum range of 3MPa. The microchannels can be easily braced by the deionized (DI) water from the syringe pump and have a pressure resistance of at least 2 MPa. When the water is pumped out, the neurotentacles can rapidly return to the membrane shape. As shown in **Fig.3D**, there is a significant difference between the probe morphology in the unfilled and filled states. These results primarily demonstrate the feasibility of our proposed method for preparing ultra-thin microchannels as well as the reliability and robustness of the microchannels.

### 2.4 Highly tunable stiffness of neurotentacles with liquid pressure

The strength of the neurotentacle is related to the pressure inside the microchannel. Quantitative tests were conducted to investigate the relationship between the maximum loaded force (F_MAX_) and the liquid pressure of the neurotentacles. The experimental setup is shown in **Fig.4A**. The neurotentacle was fixed to a motorized stage, which can determine the downward displacement (Z). A microbalance was used to test the loaded force (F_L_) on the probe. The relationships of “F_L_-Z” under different pressures are shown in **Fig. 4B**. The pressure of 0.1 MPa was used as a control, corresponding to the unfilled state and indicating that the pressure inside and outside the microchannel is equal. A tungsten wire with a diameter of 30 μm and a length of 15 mm was also used as a control. The origin of displacement (Z = 0) denotes that the probe tip had just touched the microbalance. The results show that the neurotentacle at all pressures, except for the unfilled state, exhibits similar mechanical properties. The loaded force rises rapidly as the displacement initially increases, reaching 90% of the maximum value within 200 μm (**Fig.S3**). Afterward, the force increases slowly with displacement until reaching the maximum value and then falls smoothly. Eventually, at a certain displacement, the force suddenly drops to a very small value. The states of the tested probe corresponding to this curve are shown in **Fig.4C**. It can be clearly seen that the F_MAX_ of the probe increases significantly compared to the unfilled state (0.1 MPa) and rises apparently as the pressure increases. When the pressure reaches 1 MPa, the F_MAX_ of the neurotentacle is close to that of the control tungsten wire.

**Fig 4.**
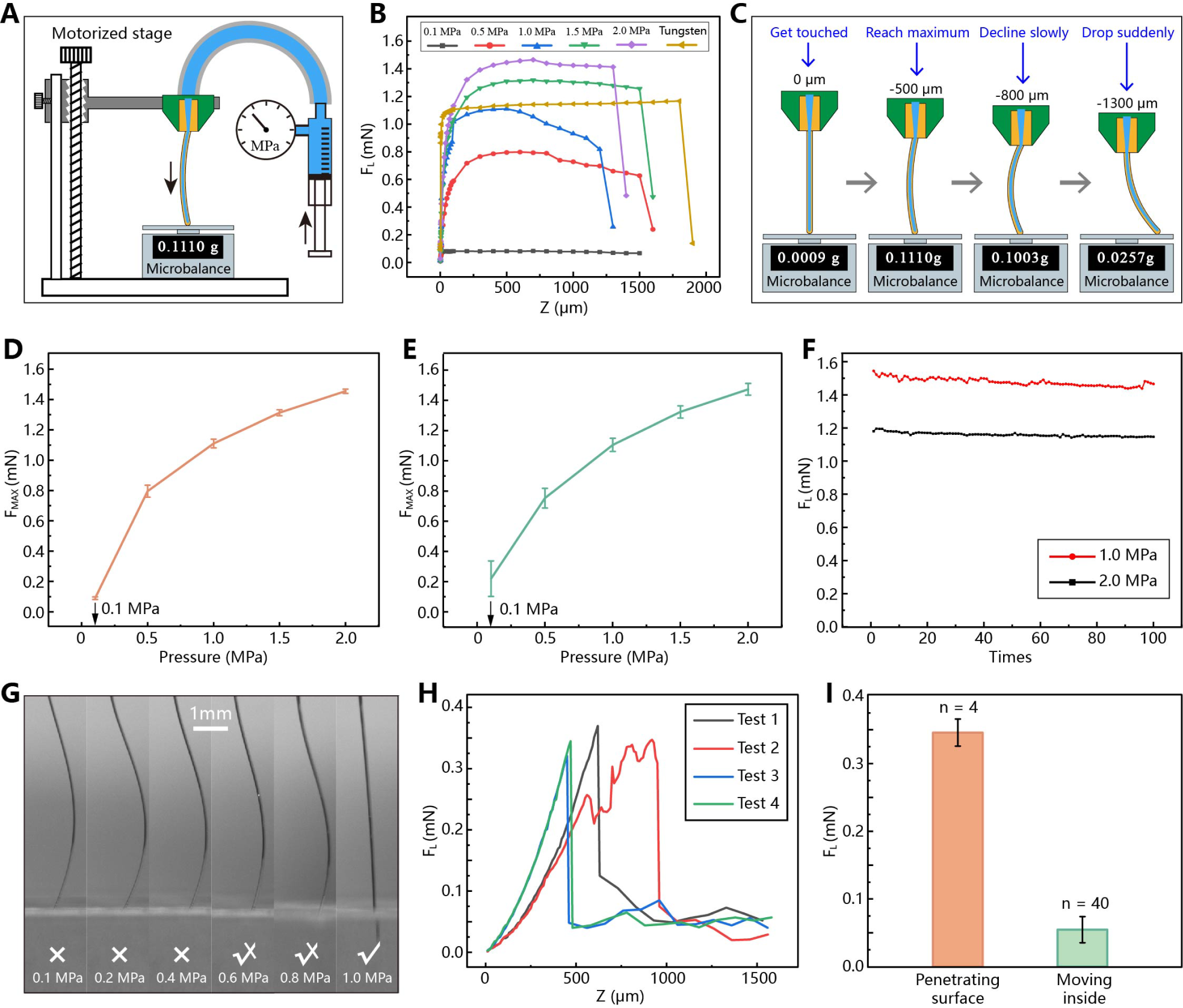
Mechanical performance of neurotentacles. (A) Experimental setup for testing neurotentacles based on a motorized stage and microbalance. (B)The relationship of loaded force (F_L_) and downward displacement (Z) for neurotentacles under different pressures and the control tungsten wire. (C) Corresponding states of the tested neurotentacles at different stages of the “F_L_-Z” curve. (D) The relationship of average FMAX and pressure (n = 10) for a neurotentacle. (E) The relationship of average FMAX and pressure for different neurotentacles (n = 4). (F) Fatigue testing of a neurotentacle at the pressure of 1 MPa and 2 MPa. (G) Testing whether neurotentacles can penetrate the brain-like gel at different pressures (0.1∼1.0 MPa). (H) The “F_L_-Z” relationships for 1-MPa neurotentacles on the brain-like gel. (I) The force required for neurotentacles to penetrate the gel and to travel inside the gel.

The "F_L_-Z" test was conducted for 10 cycles from 0.1 to 2 MPa, and the F_MAX_ at different pressures was recorded. As shown in **Fig.4D**, there is a clear positive correlation between F_MAX_ and pressure. It indicates that the strength of the neurotentacle improves significantly with internal pressure. At a pressure of 1 MPa, the F_MAX_ has risen to 1.2 mN, more than an order of magnitude above the unfilled state (80 μN). The same tests were performed with more neurotentacles and the results were highly consistent (**Fig.S4**). The average result for all the tested probes is shown in **Fig.4E**. The presented relationship between F_MAX_ and pressure is in agreement with **Fig.4D**, indicating a good mechanical consistency among different neurotentacles.

### 2.5 High robustness of neurotentacles

Fatigue tests were performed to verify the robustness of the neurotentacles under high internal pressures. The displacement required for the probe to reach F_MAX_ at a given pressure was first determined. Then, the probe was rapidly dropped from the origin (0 μm) to the position of F_MAX_, and the loaded force at that point was recorded. The experiments were repeated 100 times at 1 MPa and 2 MPa, respectively. As shown in **Fig.4F**, the same neurotentacle exhibits a negligible decrease in loaded force after 200 tests, demonstrating that the microchannel was still intact. The high robustness of the neurotentacles ensures their safety during insertion and allows them to be implanted repeatedly.

### 2.6 Critical pressure and force for inserting neurotentacles into the brain-like gel

To test whether the enhanced neurotentacle can penetrate the brain tissue and how much the critical pressure is for insertion, we performed simulated tests in the 0.6% agar gel. The experimental setup is shown in **Fig.S5**. The test pressure range is from 0.1Mpa to 1MPa. The results (**Fig.4G** and **Movie S2**) showed that the neurotentacle was unable to penetrate the gel at pressures of 0.1, 0.2, and 0.4 MPa. At pressures of 0.6 and 0.8 MPa, the neurotentacle could penetrate the gel with some probability, which may be due to the inhomogeneous surface morphology of the agar gel. When the pressure reaches 1 MPa, the neurotentacle can be inserted into the gel successfully in every test. Thus, the critical pressure for penetrating the brain-like gel should be under 1 MPa, much less than the pressure resistance (∼ 2 MPa) of the neurotentacles. This suggests that the neurotentacles are strong enough to cope with more complex situations in the brain tissue.

To further investigate the critical force for the 1-MPa neurotentacles to penetrate brain tissue, the relationship between the loaded force and downward displacement was measured on the gel, as shown in **Fig.S6**. The results of multiple tests were consistent (**Fig.4H**). The loaded force on the probes first rose and then suddenly fell with displacement. Unlike in **Fig.4B**, the loaded force rises more slowly, probably due to the deformation of the soft gel partially compensating for the increase in displacement. At about 0.35 mN, the loaded force dropped abruptly, which indicated that the probes had penetrated the surface of the gel. This further demonstrates that the neurotentacles are strong enough to penetrate the brain-like gel, as it has an MLF of 1.2 mN at 1 MPa. In addition, we found that after breaking through the surface, the force required for traveling inside the gel is extremely small (**Fig.4I**). It is only about 50μN, even lower than the MLF of unfilled neurotentacles.

### 2.7 Slight volume increase of the neurotentacle under pressure

The neurotentacle can be implanted without the use of rigid tools or temporary materials, thereby preventing a significant increase in volume during the implantation process. Upon application of pressure, the neurotentacle will change its morphology due to the expansion of the microchannel, accompanied by a slight increase in volume. The cross-sections of the neurotentacle in different states were obtained using a flip mold of PDMS (**Fig.5A** and **Fig.S7**). There are five different pressure states: 0.0 MPa (the microchannel is unopened), 0.1 MPa (the microchannel is open but not filled), 0.2 MPa, 0.5 MPa, and 1 MPa. The extracted cross-sections under different states are shown in **Fig. 5B**. At a pressure of 0.1 MPa, the area of the cross-section increases slightly compared to 0 MPa, but the overall shape remains as a sheet. At 0.2 MPa, there is a significant change in the cross-sectional morphology and the microchannel turns into an elliptical shape. At 0.5 and 1 MPa, the microchannel is getting closer to circular and the area increases slightly. We extracted the parameters of each cross-section by software and defined its maximum width (W), maximum thickness (T), circumference (C), and area (A), as shown in **Table S1**. Since Young’s modulus of PI is as high as GPa[13], the neurotentacles are unlikely to deform plastically at a pressure of MPa scale. Thus, the circumference of the probe should be relatively constant. This was used to calibrate the other dimensions of the cross-sections. The results are shown in **Table 1**.

**Fig 5.**
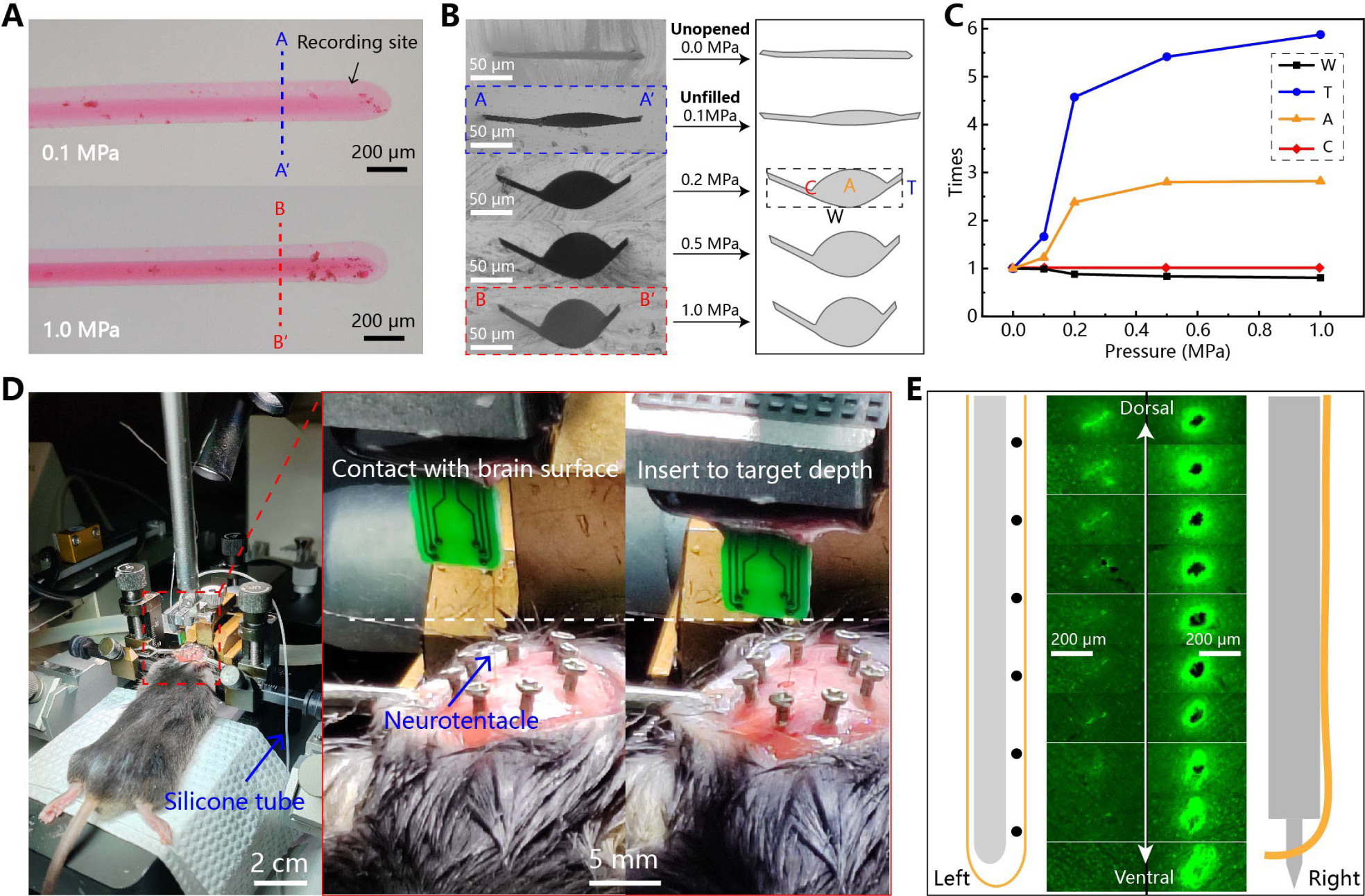
Morphology of neurotentacles and their in vivo implantation. (A) Models of neurotentacles under different pressures obtained based on the flip-mould of PDMS. (B) Cross-sections of neurotentacles at different pressures. (C) The relationship between the cross-sectional parameters of neurotentacles and the pressure. (D) Implantation of a neurotentacle in the left hemisphere of the mouse. (E) Comparison of acute implantation damage between the neurotentacle and the IFAT method

**Table 1.** Calibrated cross-sectional parameters of a neurotentacle.

| Pressure (MPa) | Width (μm) | Thickness (μm) | Area (μm<sup>2</sup>) | Circumference (μm) |
| --- | --- | --- | --- | --- |
| 0.0 | 182.4 | 9.9 | 1404.0 | 370.0 |
| 0.1 | 180.2 | 16.5 | 1724.3 | 370.0 |
| 0.2 | 160.8 | 45.3 | 3340.6 | 370.0 |
| 0.5 | 152.5 | 53.6 | 3931.1 | 370.0 |
| 1.0 | 146.9 | 58.2 | 3959.0 | 370.0 |

Defining the value at 0 MPa as 1, the trend of each cross-section parameter with pressure is shown in **Fig.5C**. As the pressure rises, the width decreases, the thickness increases and ultimately the cross-sectional area is enlarged. At a pressure of 1 MPa, although the electrode thickness increases almost five times, the final cross-sectional area only rises by a factor of about 1.8 due to the reduced width. A typical microneedle based on the IFAT method has a diameter of 100 μm[25] and a cross-sectional area of 7854 μm^2^, which ideally increases the implantation volume by at least 5.6 times. This suggests that the neurotentacle can reduce implantation damage by nearly 60% compared to traditional implantation methods for flexible electrodes. Furthermore, the implantation damage might be even lower if inserting the neurotentacle with varied pressure, according to **Fig.4I**.

### 2.8 Low invasiveness of neurotentacles

To evaluate the acute damage, a neurotentacle was implanted into the zona incerta (ZI) in the left hemisphere of the mouse. As shown in **Fig.5D**, the neurotentacle was carried on a stereotaxic device for the implantation as conveniently as a rigid probe. Saline was used as the internal liquid. At a pressure of 1 MPa, the neurotentacle could be easily inserted into the brain tissue. The insertion speed was 100 μm/s and the insertion depth was 4.4 mm. Then the saline in the neurotentacle was pumped out to recover its flexibility. In the right hemisphere, a conventional PI flexible probe (**Fig.S8**) was implanted based on the IFAT method as a control. The flexible electrode for the control was 180 μm wide and 6 μm thick. A tungsten needle with a diameter of 100 μm was used to insert it (**Fig.S8**), using the same insertion parameters. The tungsten needle was then withdrawn, leaving only the flexible probe inside the brain (**Fig.S9**).

Next, we performed immunostaining characterizations with neuronal nuclear protein (NeuN) to assess acute implantation damage between the neurotentacle and the IFAT method. Since the recording sites of the electrodes were placed at the tip, the brain slices were randomly selected at the appropriate depth and sequenced from dorsal to ventral (a distance of approximately 0 ∼ 600 μm from the tip of the neurotentacle). It is evident that the neurotentacle resulted in significantly less insertion damage than the IFAT method. In the right hemisphere, the flexible electrode introduced by the tungsten needle created a hollow with a diameter of approximately 50 μm (**Fig.5E**). However, in the left hemisphere, only minimal residual evidence remained in the region where the neurotentacle had been implanted. This result indicated that the neurotentacle caused minimal acute damage compared to conventional flexible probe implantation methods. This could potentially reduce the immune response of the surrounding tissue after implantation and enhance the signal quality of long-term electrophysiological recordings.

To further elucidate the effects of long-term retention of electrodes on tissue damage and its immune response, a mouse was implanted with neurotentacles and its brain tissue was obtained by perfusion five weeks later. Following the preparation of frozen sections of the brain, those sections in proximity to the probe tips were subjected to NeuN and glial fibrillary acidic protein (GFAP) immunofluorescence histochemistry, respectively, to determine the extent of brain tissue damage and the immune response. In comparison to the right hemisphere, where no electrodes were implanted, no discernible indications of immune response were observed in the left hemisphere, where the neurotentacle had been implanted (**Fig. S10**). Furthermore, the trace of the probe path was nearly undetectable. These results indicated that the neurotentacle exerted minimal influence on the surrounding neurons and immune response following chorionic implantation.

### 2.9 Stable electrochemical properties of neurotentacle under varied pressures

Furthermore, the electrochemical characteristics of the encapsulated neurotentacles were tested. In special, the frequency–impedance spectroscopy and cyclic voltammetry (CV) curves of the electrodes were recorded before and after modification with Poly(3,4-ethylenedioxythiophene) (PEDOT). The results are shown in **Fig.6**.

**Fig 6.**
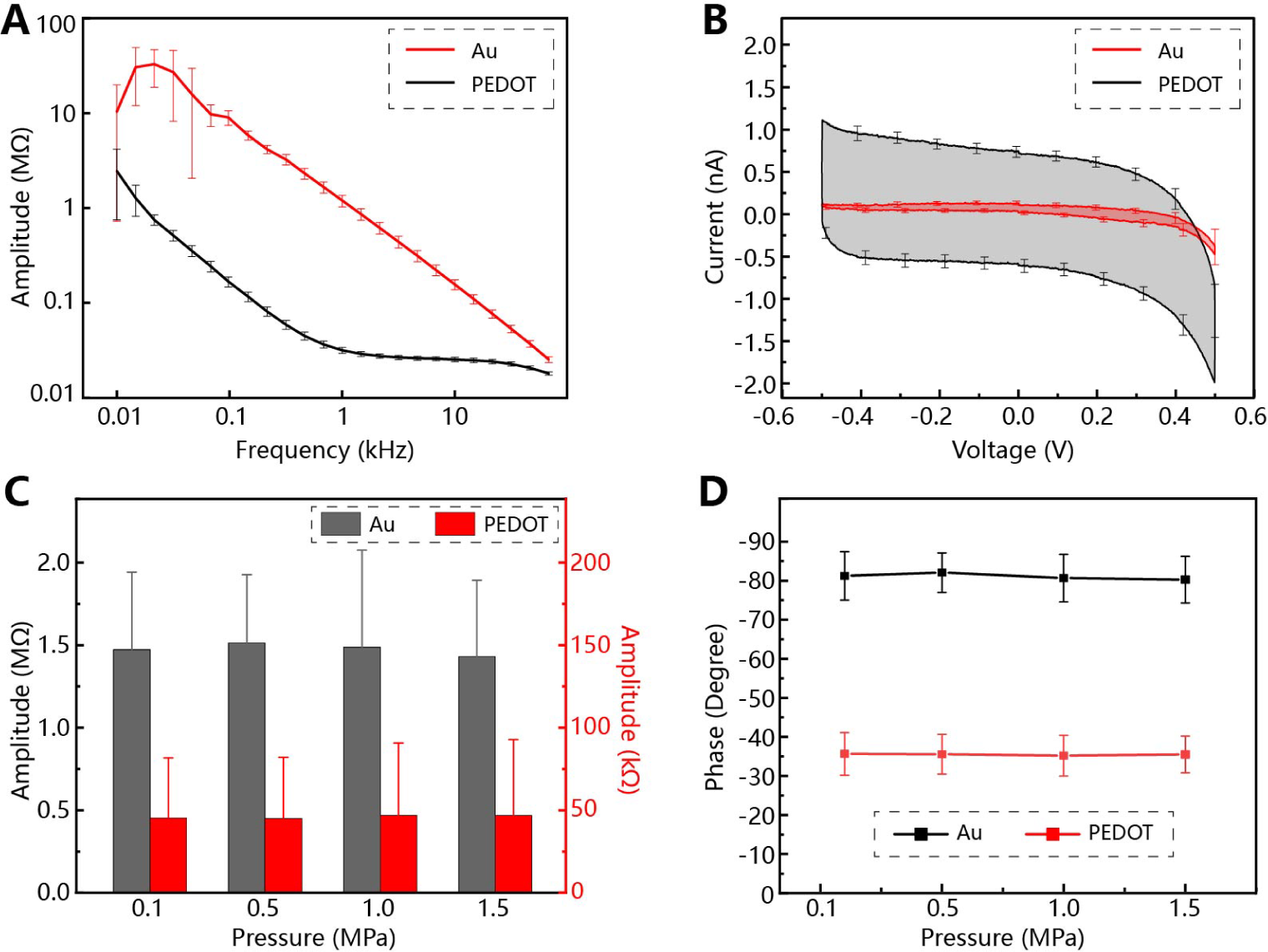
Electrochemical characterization of the recording sites on neurotentacles before and after modification with PEDOT. (A) Frequency-impedance spectra (n=20). (B) CV curves (n=20). (C) Effect of pressure on impedance at 1kHz (n=14). (D) Effect of pressure on phase at 1kHz (n=14).

The average impedance amplitude decreased significantly after the modification (**Fig.6A**). At 1kHz, the amplitude dropped from 1.36 ± 0.35 MΩ (mean ± s.e.m.) to 33.5 ± 10.8 KΩ. The CV curves show that the charge storage capacity of the modified electrodes was also significantly increased (**Fig.6B**). There is no significant difference between the electrochemical performance of the neurotentacles and that of normal PI probes[6]. We also tested the effect of internal pressure on electrode impedance. **Fig.6C** and **Fig.6D** show that the electrode impedance does not change with the pressure in the microchannel, either before or after modification with PEDOT. These results demonstrate that the electrical properties of neurotentacles are stable enough to withstand the microchannel volume changes induced by pressure variations.

### 2.10 Stable recording of neurotentacles in vivo

To evaluate the stability of the neurotentacles to obtain the electrophysiological signals of neurons, the hippocampus CA1 region, which is the most classic area in the field of electrophysiological recording, was selected as the brain area for electrode implantation in this stage. After being implanted with neurotentacles for two months, spikes with high SNRs were acquired in all the mice. A total of 26 valid channels were identified across the four probes, with in vitro impedance measurements below 100KΩ at 1 kHz. The recorded continuous data of these channels on day 16 is shown in **Fig.7A**. There were 23 channels that had acquired spikes with a yield of 88.5%. A total of 75 units were sorted out from these channels, as shown in **Fig.7B**. The spike autocorrelograms and interspike intervals of four example units from channel-8 of mouse-2 are presented in **Fig.7C**.

**Fig 7.**
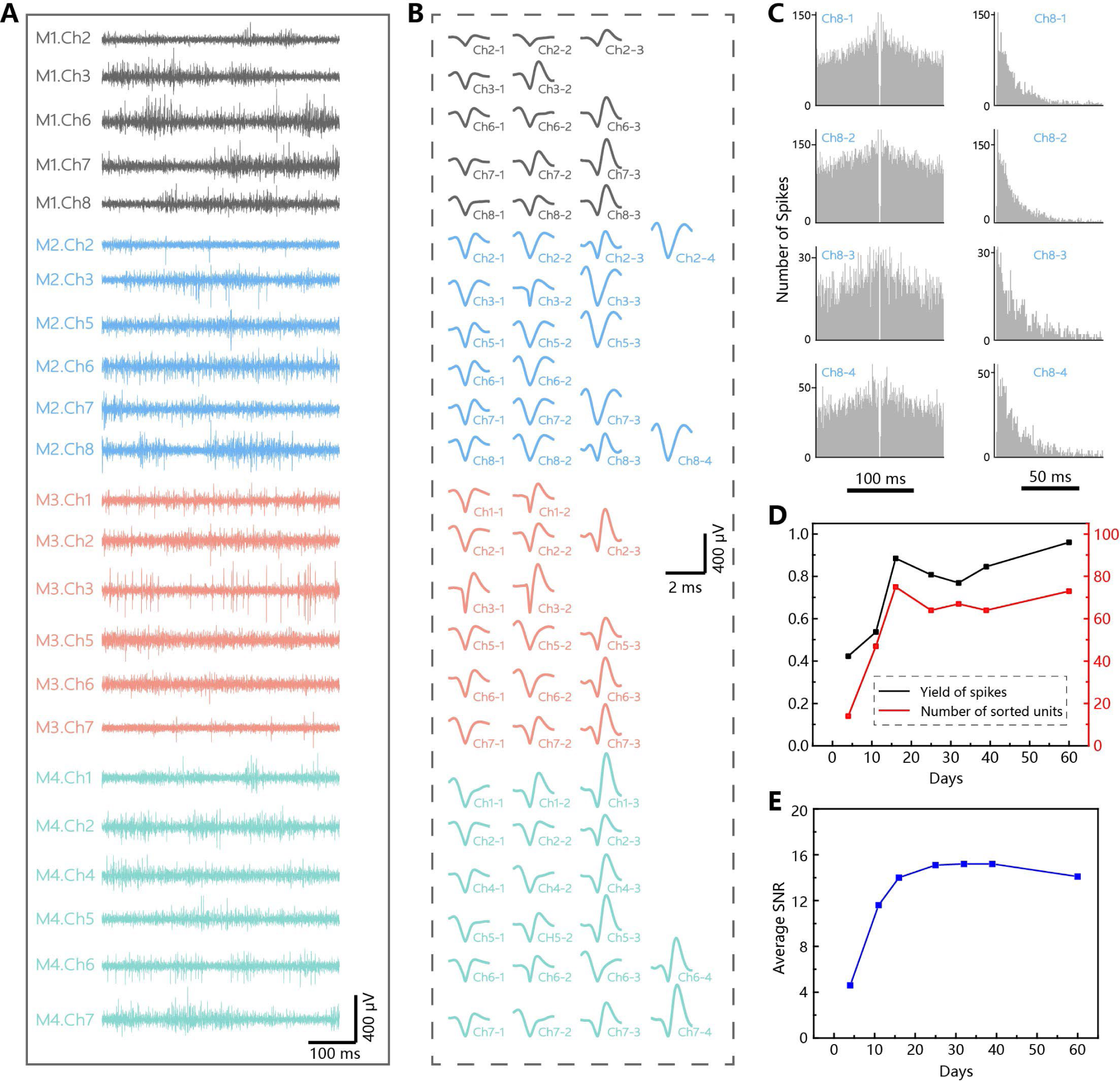
In vivo recordings of neurotentacles on mice. (A) Continuous data of 23 channels from 4 mice where spikes had been acquired on the 16^th^ day. “M1.Ch2” indicates channel 2 on mouse #1. (B) Sorted units from the corresponding channels in (A). “Ch2-1” indicates the unit #1 from channel 2. (C) The spike autocorrelograms and interspike intervals of four example units from M2-Ch8. (D) Evolution of spikes yield and number of sorted units over two months. (E) Evolution of average SNR over two months.

The evolution of the yield of all the valid channels over a two-month period is shown in **Fig.7D**. During the initial two-week period, the spikes yield increased from 42.3% to 88.4%, accompanied by an increase in the number of units sorted from 14 to 75. The yield was maintained at this high level for the next five weeks. On the 60^th^ day following implantation of the neurotentacles, the spike yield reached a maximum of 96%. The average SNR was calculated for all the channels that had acquired spikes on the same day. As illustrated in **Fig.7E**, the stable chronic recordings of the average SNR increased from 4.1 to 14 within the initial two-week period and remained above this value thereafter. These data demonstrated the excellent in vivo performance of neurotentacles, implying their potential for long-term recording.

## 3 Discussion

### 3.1 Advantages of using the FMAX to assess the strength of neurotentacles

The Euler buckling load (F_B_) is often used to characterize the buckling strength of neural probes. It can be denoted as[35, 36]:

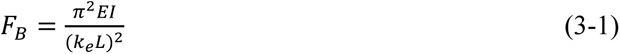

where *E* is Young’s modulus, *L* is the effective length, *k_e_* refers to the effective length coefficient that depends on the boundary conditions and is on the order of 1, and *I* is the moment of inertia and it is related to the shape and size of the section. However, in this paper, we use F_MAX_ rather than F_B_ to describe the variable strength of the neurotentacle. F_MAX_ and F_B_ can be both obtained from the “FL-Z” curve. For the test procedure in **Fig.2**, if the electrode tip is fixed so that it ends up bending instead of moving away, then F_B_ corresponds to the snap point. There should be a positive correlation between F_MAX_ and F_B_. Thus, it is feasible to characterize the strength of the neurotentacle in terms of F_MAX_, which is equally determined by the parameters in Equation 3-1. In addition, the measurement of F_MAX_ does not involve any buckling of the device. This allows the same neurotentacle to be tested repeatedly under different conditions, with no mutual influence among the results and no risk of breaking the electrode traces.

### 3.2 Factors affecting the maximum strength of neurotentacles

As depicted in **Fig.4D**, there is a clear positive correlation between the strength of the neurotentacle and the applied liquid pressure. However, the strength increase is gradually slowing down as the pressure rises. We speculate that this is because the pressure regulates the strength primarily by causing changes in the probe footprint and secondarily by altering Young’s modulus. For the neurotentacle, both Young’s modulus and the footprint are variable with pressure. As shown in **Fig.1**, the probe size increases rapidly when the pressure begins to rise, which may result in faster growth of strength at first in **Fig.2**. After that the probe size tends to stabilize and the strength increases more slowly with the equivalent Young’s modulus. Although increasing the liquid pressure can theoretically keep improving the strength of the probe, actually exceeding a pressure of 3 MPa will challenge the sealing of the whole liquid-connected chamber. What is more critical in determining the upper limit of the strength is the footprint of the probe, i.e. the size of the microchannel.

### 3.3 Ultra-low invasiveness of neurotentacles

Through in vitro and in vivo experiments, the neurotentacle was demonstrated with low invasiveness. Actually, the invasiveness can be further reduced. We found that the F_MAX_ of neurotentacle at 1MPa is close to 1.2 mN (**Fig.4B**), but there needs only about 0.43mN to penetrate the brain-like gel (**Fig.4I**). In addition, the applied pressure can be increased to more than 2MPa. These results indicate that there is still room to reduce the footprint of the neurotentacle, thus reducing the insertion damage. It requires a proper compromise between the size and the maximum strength of the probe.

Moreover, the force required for traveling inside the brain-like gel is less than 100μN (**Fig.4B**). This suggests that it may be possible to penetrate the tissue with a higher pressure but move to the target depth with a lower pressure. We have validated it in the brain-like gel, as shown in **Fig.S11** and **Movie S3**. The neurotentacle penetrated the gel at 1 MPa and then continued to be inserted at 0.1 MPa This provides a possibility to achieve ultra-low invasive implantation for neurotentacles.

## 4 Materials and Methods

### 4.1 Dilution of PI2611

The dilution for PI is referred to the previous work of Liu et al[34]. The procedures are as follows: (1) Mix the PI2611 prepolymer with the NMP solution at a mass ratio of 4:1. (2) Use a magnetic stirrer to make the mixture well blended. (3) Remove the air bubbles from the mixture in a vacuum environment. (4) Obtain the diluted PI2611 prepolymer.

### 4.2 Fabrication of neurotentacle

The processes for manufacturing the neurotentacles are shown in **Fig. 2A-2B** and are described as follows: (1) Spin-coating (2000 r/s) diluted PI2611 on the silicon wafer and baking (300 ℃) to form the bottom PI layer. (2) Patterning the positive photoresist (AZ4620) on the bottom PI to define the microchannel regions. (3) Treating the surface with reactive ion etching (RIE, 90 s) to create a strongly-adhesive region around the microchannels. (4) Patterning the negative photoresist (AR4340) with the microchannels being uncovered. (5) Treating the microchannels with hydrophobic reagents (1H,1H,2H,2H-tridecafluorooctyl) to further reduce the adhesion. (6) Spin-coating (5000 r/s) diluted PI2611 and baking (300 ℃) to form the middle PI. (7) Treating the middle PI with RIE (O_2_, 30 s) to increase its adhesion. (8) Patterning the photoresist (AR4340), depositing the composite metal layers (Cr/Au/Cr), and lifting off the photoresist to obtain the metal traces and electrodes. (9) Spin-coating (4000 r/s) undiluted PI2611 and baking (300 ℃) to form the top PI. (10) Patterning the photoresist (AZ4620), RIE etching the PI, and lifting off the photoresist to define pads, recording sites, and probe profiles. (11) Etching the upper Cr layer on the pads and recording sites and annealing the wafer on a hot plate (350 ℃) for 30 minutes. (12) Soaking the wafer in deionized water and using a tweezer to release the neurotentacles from the substrate.

### 4.3 Encapsulation of neurotentacle

The procedures for encapsulating the neurotentacle with water and electrical connectors are shown in **Fig.S2**, and they are described as follows: (1) Opening the microchannel from the rear end with a syringe needle (outer diameter: 0.18 mm). (2) Inserting the syringe needle into the microchannel until it closely fits the inner PI wall. (3) Sealing the joint with UV glue. (4) Connecting the balloon syringe pump to the syringe needle and inject liquid to open the whole microchannel. (5) Cutting off the syringe with only the stainless-steel needle left. (6) Electrically connecting the neurotentacle to the PCB with gold ball bonding[6, 25]. (7) Attaching a silicone tube (inner diameter: 0.3 mm, outer diameter: 0.8 mm) to the needle and sealing the joint with UV glue. (8) Connecting a syringe needle (outer diameter: 0.18 mm) to the other end of the silicone tube and sealing the joint. (9) Soldering the connector (pin pitch: 1.27 mm) to the PCB. (10) Sealing the whole PCB with UV glue to form a waterproof and insulating encapsulation.

### 4.4 Mechanical testing of neurotentacle

During testing the "load force-displacement-pressure" relationship for neurotentacles, a motorized stage was used to carry the probe up and down with a minimum step distance of 1μm, a microbalance was used to measure the load force with a minimum readability of 1μN, and the hydrometer on the syringe pump was used to monitor the internal pressure with a minimum scale of 0.1 MPa.

### 4.5 Morphological testing of neurotentacle

The cross-sections of the probe under different hydraulic pressures were acquired based on the flip-mould of PDMS. The procedures are shown in **Fig.S7** and are described as follows: (1) Placing a mold on the hot plate (20 ℃) and holding the neurotentacle vertically inside the mold. (2) Filling the mold with PDMS (Sylgard 184, Dow Corning, with the weight ratio of prepolymer to curing agent of 10:1) until the neurotentacle is submerged about 5 mm. (3) Curing PDMS at 120 °C for 10 minutes. (4) Removing the probe and taking the cured PDMS out of the mold. (5) Cutting off the PDMS to expose the cross-section at the depth of the recording sites. (6) Acquiring the cross-section with an optical microscope. (7) Obtaining the cross-sectional outline of the microchannel and calculating the dimensional parameters with software (Matlab).

### 4.6 Electrochemical characterization of neurotentacle

An electrochemical workstation (CHI660D) was used to perform frequency–impedance spectroscopy, cyclic voltammetry (CV), and modification, with the microelectrodes as the working electrodes and a Pt electrode as the counter electrode. The impedance and CV were measured in PBS solution. Poly(3,4-ethylenedioxythiophene) (PEDOT) was deposited in an aqueous solution consisting of 0.02 M EDOT monomer and 0.1 M TsONa (sodium p-toluene sulfonate) electrolyte using the constant current method with a current magnitude of 6.35 nA for 30 s per electrode.

### 4.7 Animals

The surgical procedure was conducted on male mice of the C57BL/6J strain, which were 2 months old at the commencement of the experiments. Mice were housed in a specific pathogen-free (SPF) barrier environment with a 12-hour light/dark cycle control at 22 ± 1°C and with food and water ad libitum. Animals were implanted with chronic flexible neural microelectrodes and single-housed. Chronic electrophysiological recording experiments were conducted during the light phase. Prior to the commencement of the electrophysiological recording experiments, the animals were habituated to the experimenter by handling for a minimum of 3 consecutive days (10 minutes per day). Animal care and use were under the guidelines approved by the Animal Care and Use Committee in Institute of Psychology, Chinese Academy of Sciences.

### 4.8 Surgical implantation of flexible neural microelectrodes

Mice were initially anesthetized using isoflurane (induction: 5.0%, maintenance: 1.0%), before being secured within a stereotaxic frame (RWD Life Science, China) and eye ointment was subsequently applied. To evaluate the acute damage, a flexible probe introduced by the tungsten needle and a neurotentacle were implanted in either the left or the right side of the ZI (coordinates: 1.91 mm posterior, ±1.20 mm lateral, and 4.20-4.45 mm ventral from Bregma). To assess the quality of in vivo recordings, the neurotentacles were implanted in the CA1(coordinates: 1.82 mm posterior, 1.50 mm lateral, and 1.20-1.40 mm ventral from Bregma). Holes were drilled above the target brain regions. A tungsten needle guided flexible probe was implanted using the IFAT method with an insertion speed of 100 μm/s. Then the tungsten needle was withdrawn at a speed of more than 1000 μm/s, leaving only the flexible probe in the brain (**Fig.S9**). The neurotentacles were carried on a stereotaxic device for implantation (**Fig.4D**). The pressure inside the probe during insertion was 1MPa, and the insertion speed was also 100 μm/s. After reaching the specified depth, release the inside pressure to 0.1Mpa, and then cut the silicone tube at the headstage. The two implanted probes were fixed to the skull with miniature skull screws, cyanoacrylate glue, and blended dental cement.

### 4.9 Chronic electrophysiological recordings and data analysis

Mice were allowed to recover for 4 days after surgery and then acclimated to the headstage and cables for several days before electrical recording began. To explore CA1 activity, subject mice were placed in the open-field apparatus and allowed to acclimate for 10 min before starting in vivo recordings. Wideband electrophysiological data were recorded with a sampling rate of 40 kHz using the OmniPlex Neural Recording Data Acquisition System (Plexon Inc, USA). Subsequently, the raw wide-band data were preprocessed using the Offline Sorter software (Plexon Inc, USA) and further analyzed through NeuroExplorer (Nex Technologies, USA) in combination with custom-written MATLAB scripts. To detect the spiking activity of single units, a standard fixed threshold method was employed, with the amplitude threshold set at 3 standard deviations. The units were then automatically separated by principal component analysis and cluster analysis.

The signal-to-noise ratio (SNR) tool was calculated by the Offline Sorter software using the following formula:

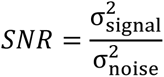

where 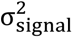 is the variance of the signal and 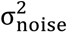 is the variance of the noise. Since there are sorted units in the spike channels, 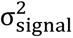 is defined as all sorted spikes, and 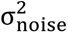 is defined as the data between all spikes.

### 4.10 Immunohistochemistry

Mice were anesthetized with isoflurane and perfused sequentially with phosphate-buffered saline (PBS) followed by 4% paraformaldehyde (PFA). For post-fixation, brains were incubated in PBS containing 30% sucrose until they sank to the bottom. Cryostat sections (100 mm perpendicular to the probes) were obtained using a cryostat (Leica CM3050S). Sections were washed in PBS 3 times (10 minutes each time) and then incubated with blocking solution (PBS containing 10% goat serum and 0.5% Triton X-100) for 2 hours at room temperature, and then treated with primary antibodies diluted with blocking solution overnight at 4℃. The following primary antibodies were used: Rat anti-NeuN (1:500; Oasis-Biofarm, OB-PRT013), and Rabbit anti-GFAP (1:500; Oasis-Biofarm, OB-PRB005). After three washes in PBS (10 minutes each), sections were then incubated with species-specific fluorophore-conjugated secondary antibodies (1:1000, goat anti rat 488nm, Oasis-Biofarm, RT488 or 1:1000, goat anti-rabbit Alexa Fluor 546, Invitrogen, A-11010) in PBS for 2 hours at room temperature. After washing in PBS 3 times (10 minutes each time), sections were stained with DAPI, washed with PBS, and transferred to microscope slides. Sections were imaged using a Laica DMi8 fluorescence microscope (10× objective lens).

## Funding

National Key R&D Program of China (2022YFF1202303)

National Key R&D Program of China (2023YFF1203702)

National Natural Science Foundation of China (62071447)

National Natural Science Foundation of China (32071028)

## Author contributions

Device design and preparation: Y.W., Y.C.

Device encapsulation: Y.W., X.Y.

Design of experiments: Y.W., X.X., W.P., J.L., Y.W.

In vitro test: Y.W.

Animal experiment: X.X., R.T.

Electrophysiological data recording: Y.W., X.X.

Electrophysiological data processing: X.X.

Histological experiments: X.X.

Writing—original draft: Y.W., X.X.

Writing—review & editing: W.P., J.L.

## Data and materials availability

All data are available in the main text or the supplementary materials.

## This PDF file includes

Figs. S1 to S11

Tables S1

Movies S1 to S3

## Supplementary Materials

**Fig. S1.**
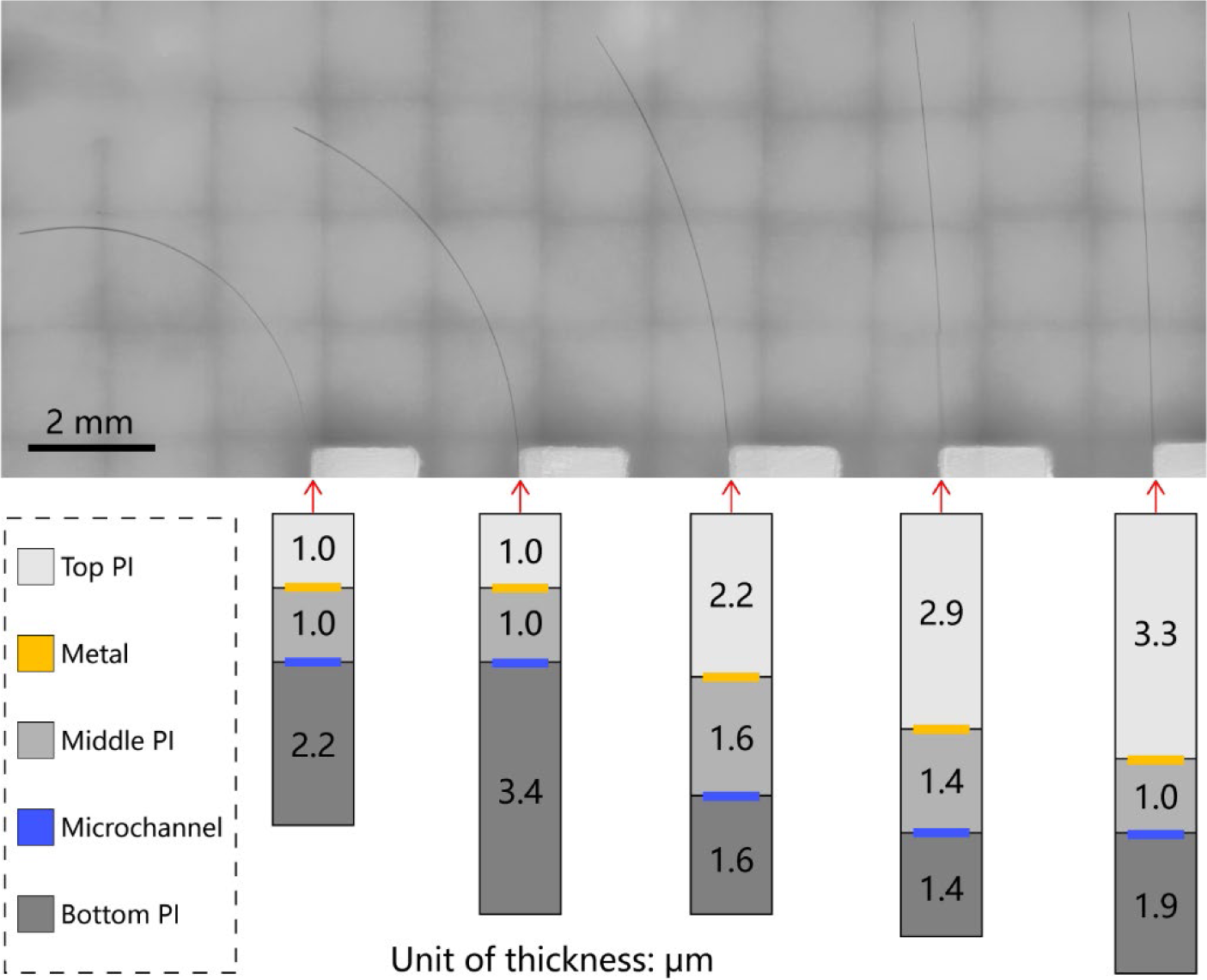
Self-bending of the neurotentacles due to uneven PI thickness distributions above and below the metal layer

**Fig. S2.**
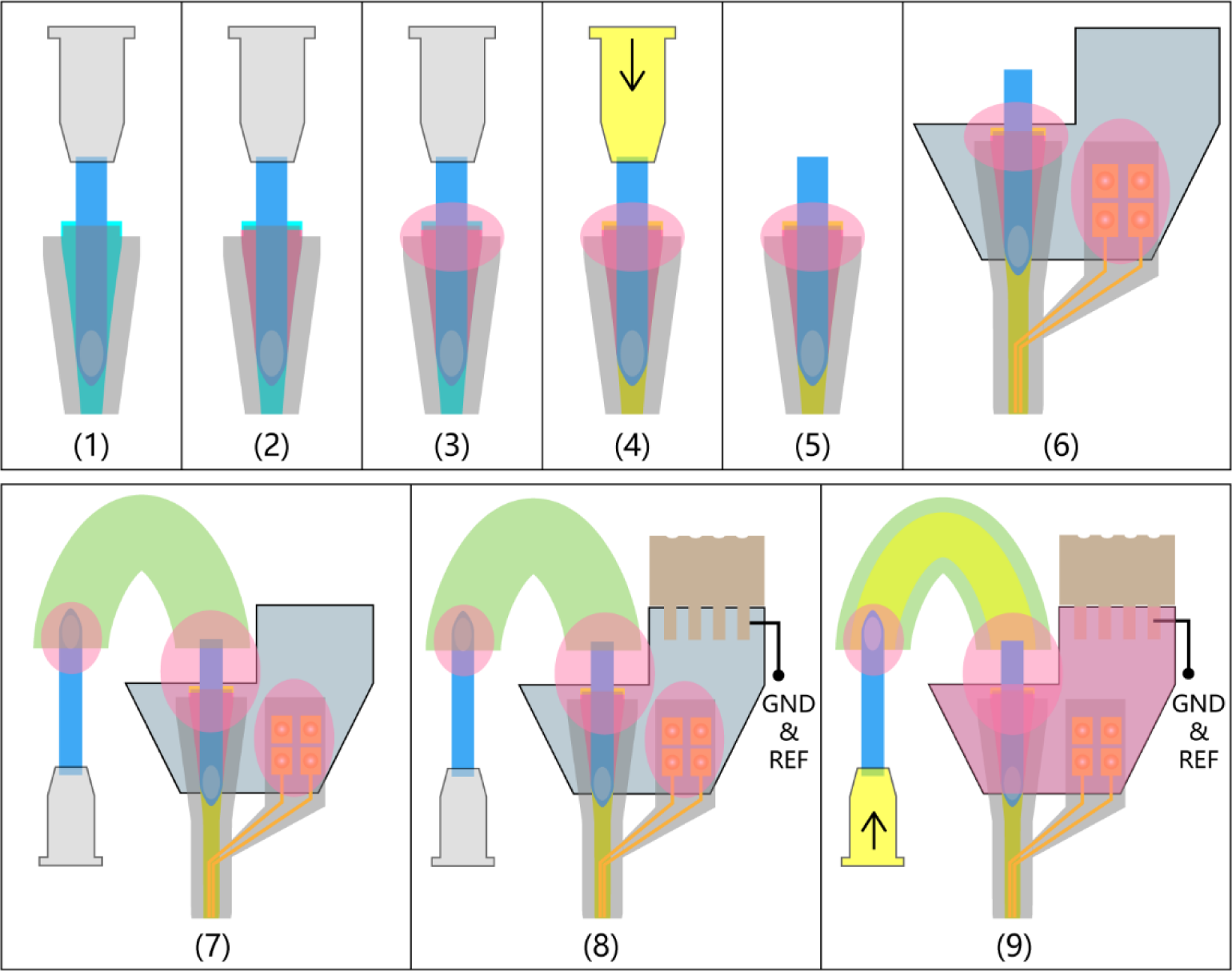
Procedures of encapsulating the neurotentacle with the liquid pathway and electrical connector.

**Fig. S3.**
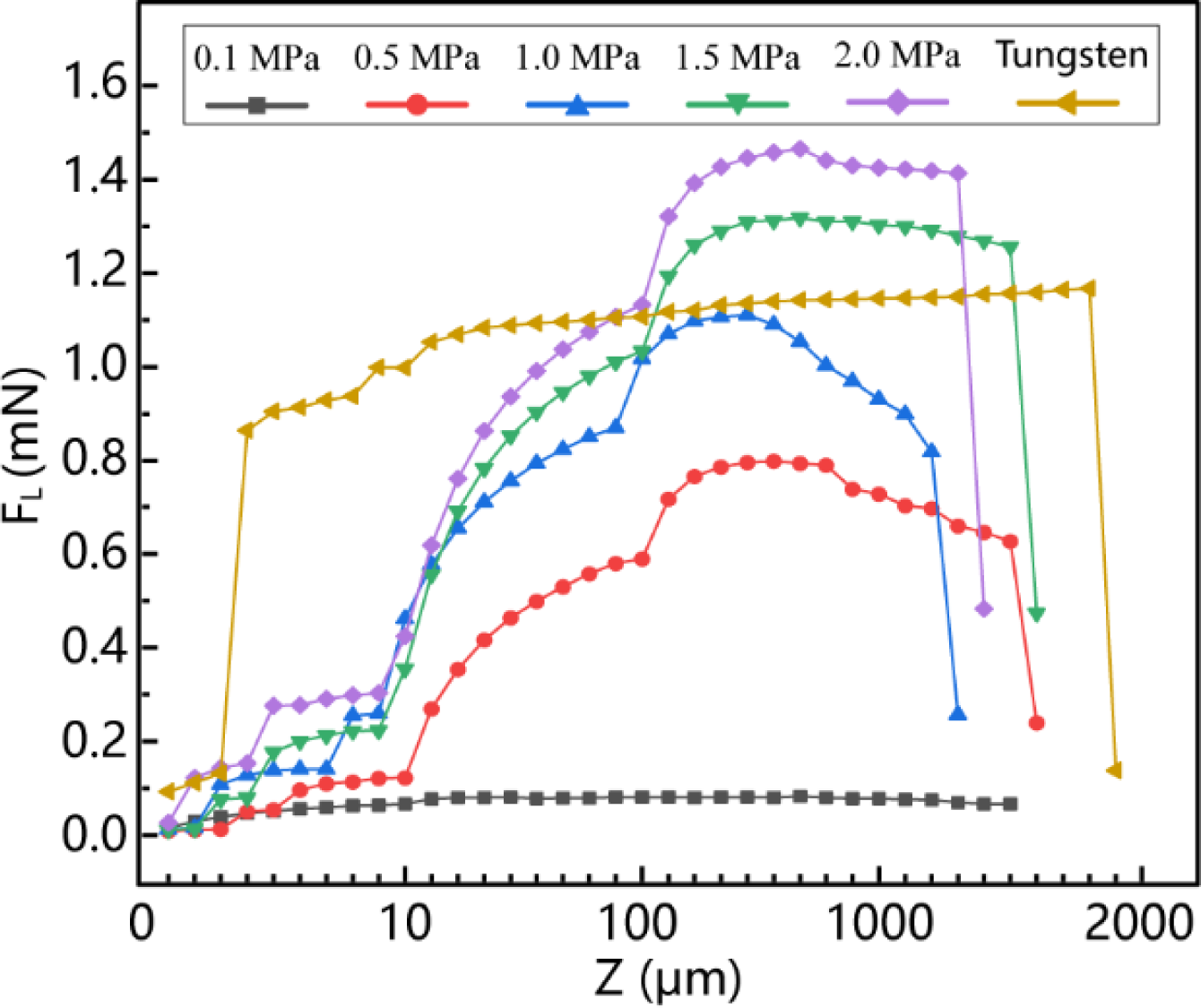
The relationship of loaded force (FL) and downward displacement (Z) for neurotentacles under different pressures and the control tungsten wire (nonlinear transverse coordinates).

**Fig. S4.**
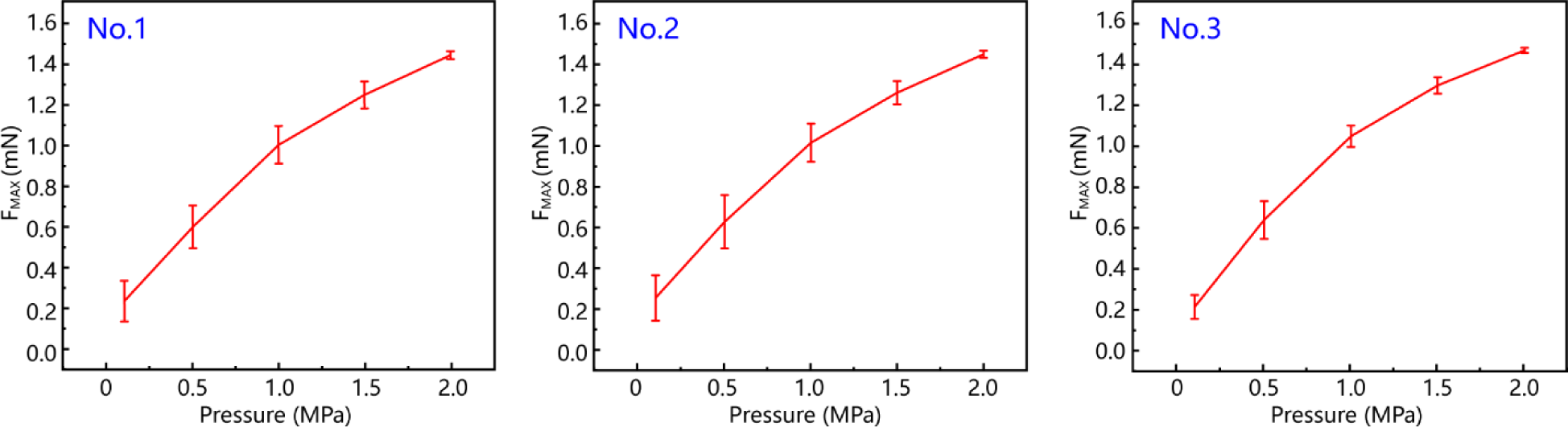
The relationships of average FMAX and pressure (n = 5) for another three neurotentacles.

**Fig. S5.**
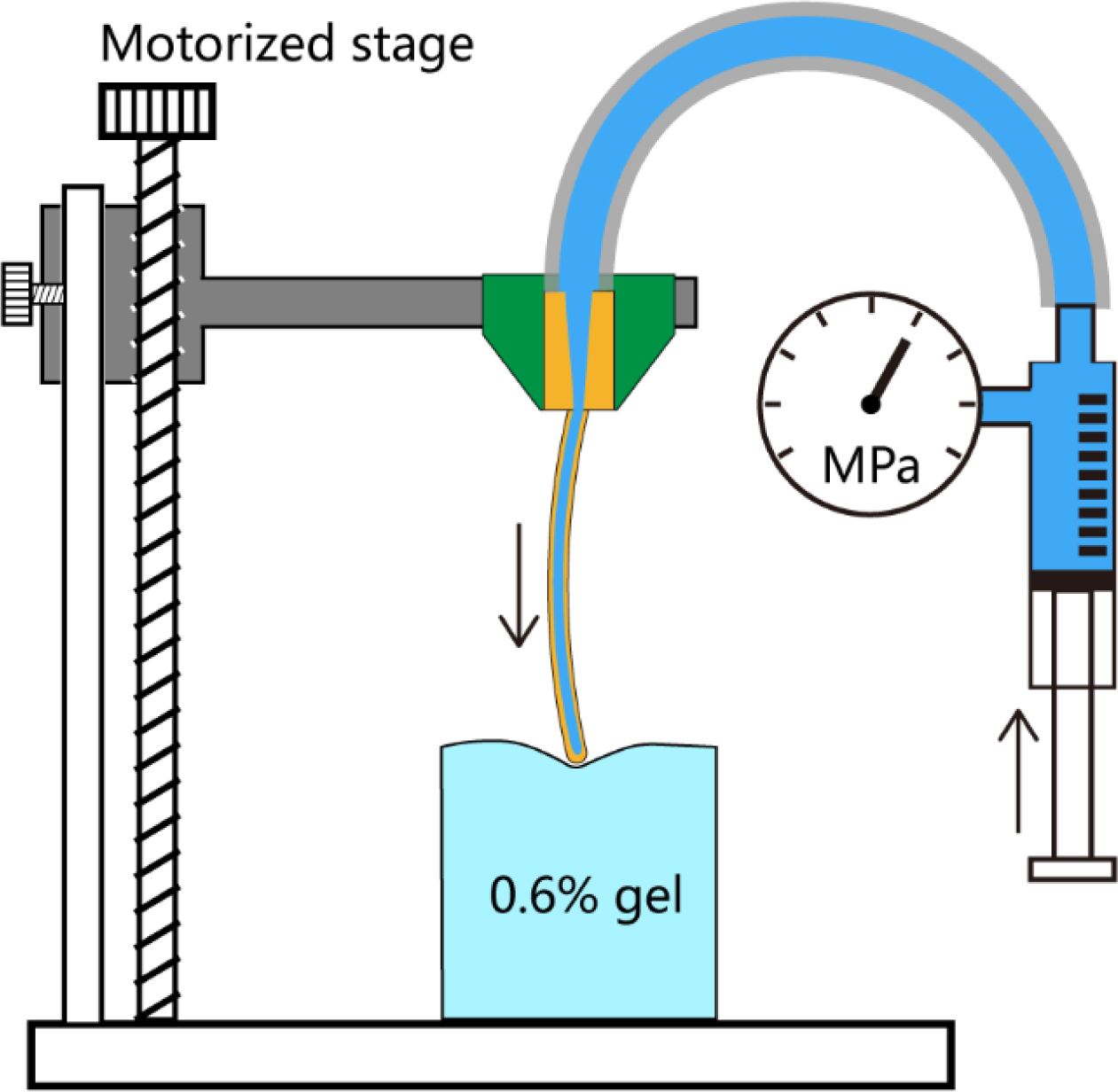
Experimental setup for testing the critical pressure of neurotentacles on the gel.

**Fig. S6.**
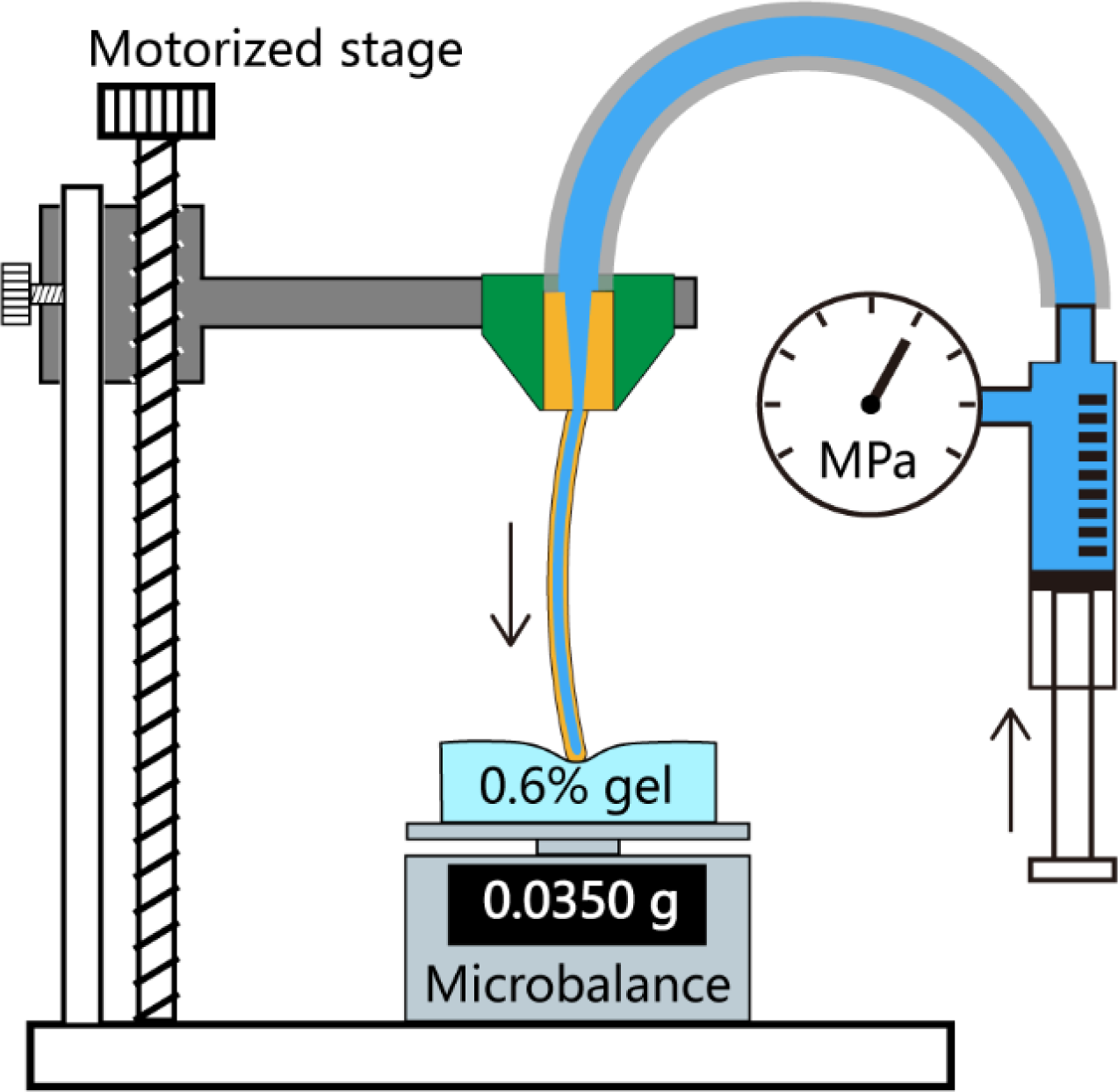
Experimental setup for testing the critical force needed for neurotentacles to penetrate the gel

**Fig. S7.**
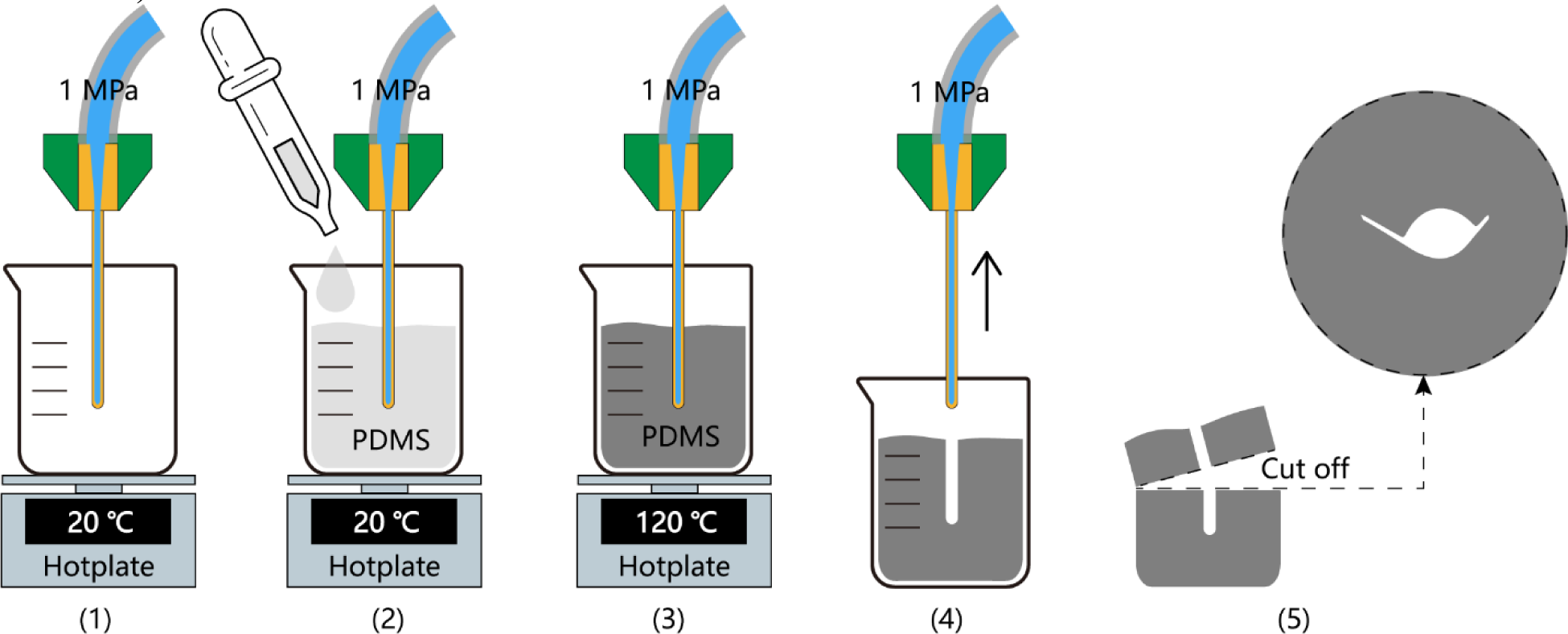
Procedures for acquiring cross-sections of neurotentacles under different pressures (1MPa for example) based on the flip-mould of PDMS

**Fig. S8.**
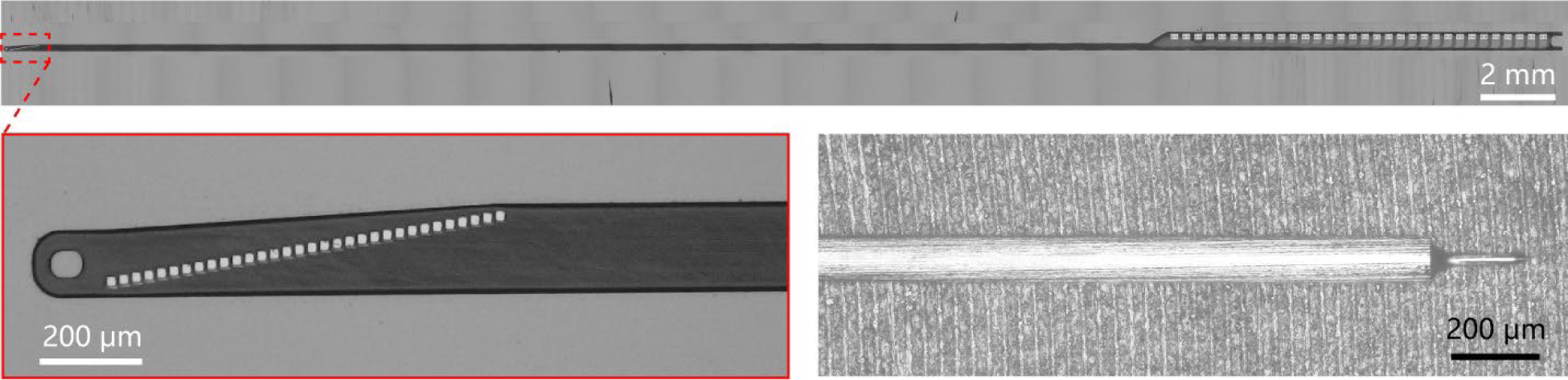
The conventional flexible probe and rigid microneedle used in the IFAT method for implantation in the right hemisphere of the mouse

**Fig. S9.**
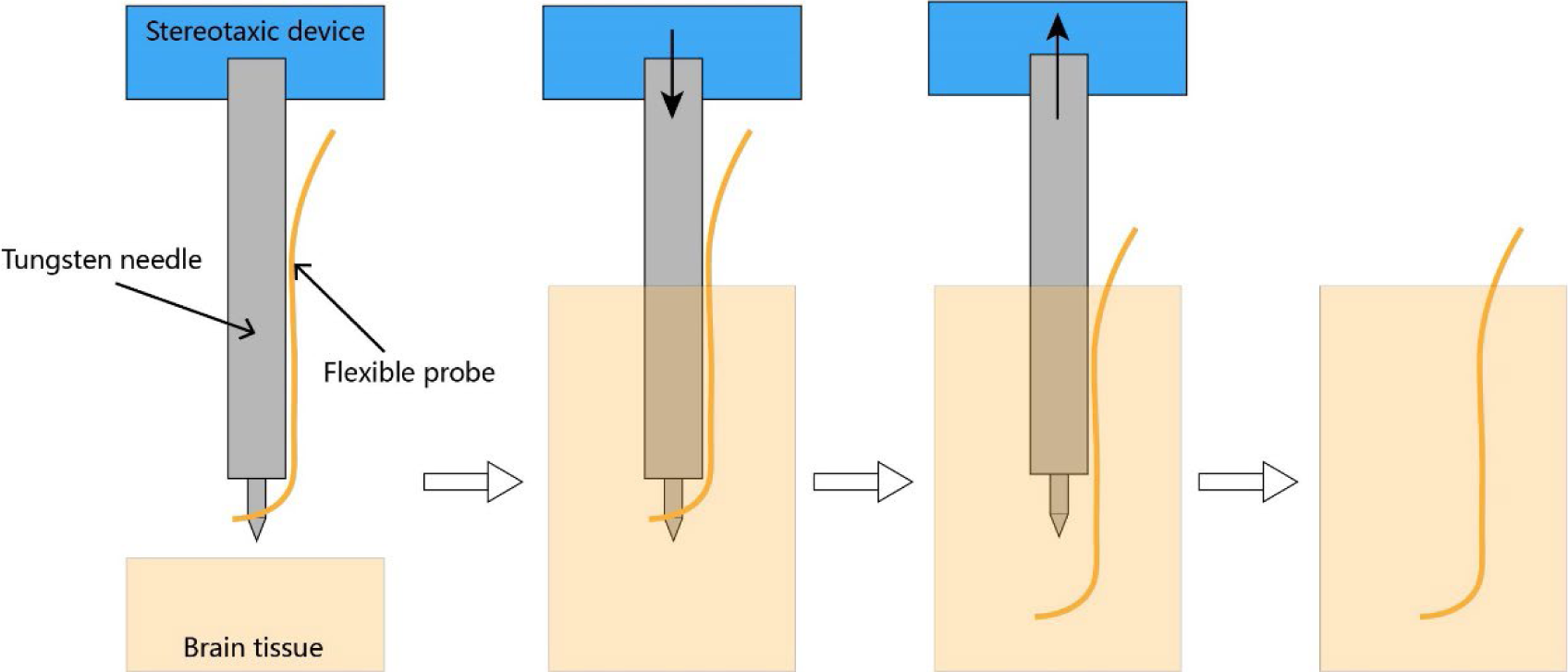
Schematic diagram of the implantation procedures for the conventional IFAT method

**Fig. S10.**
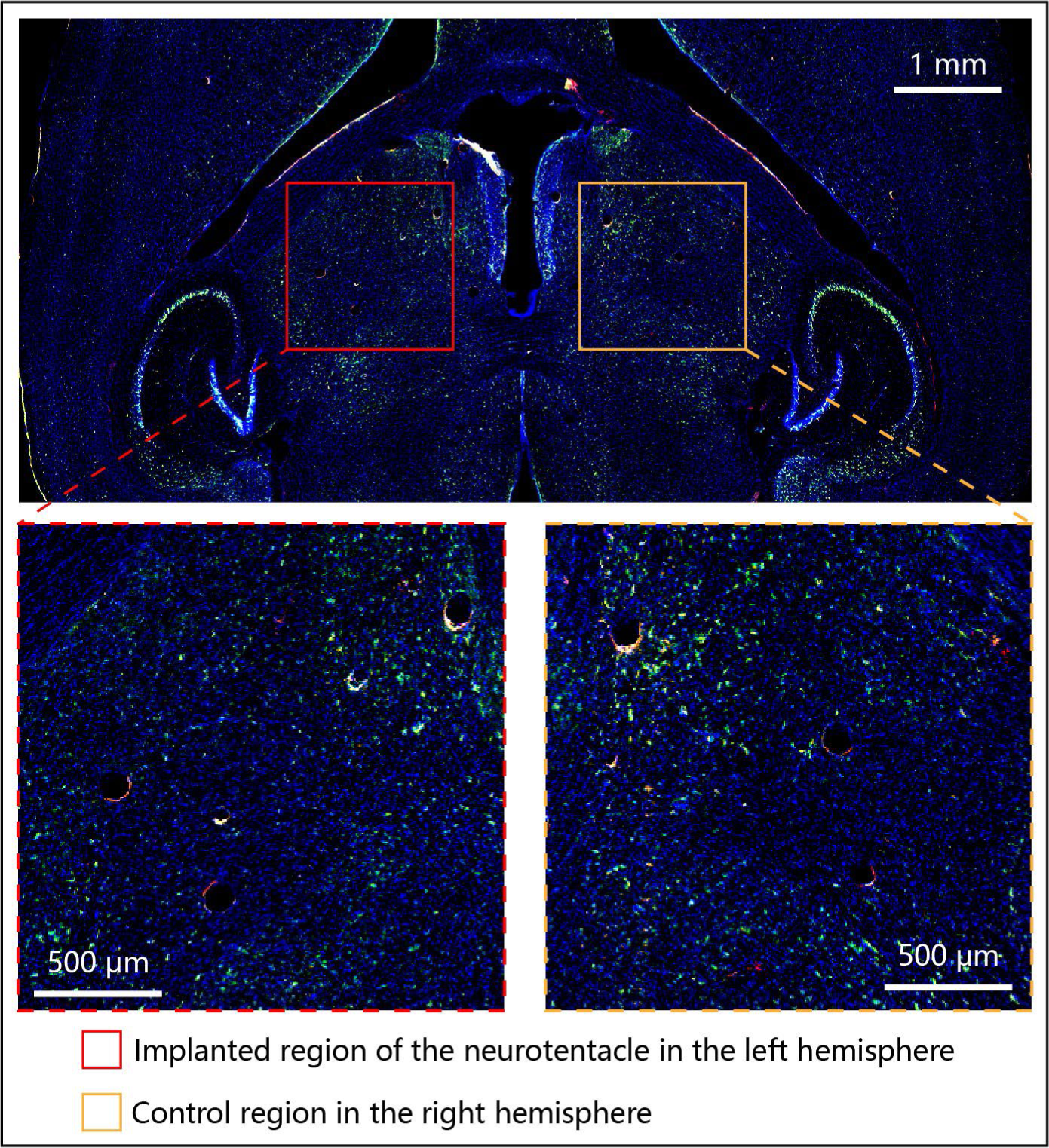
Histological sections and immunostaining of mice after implantation with a neurotentacle for five weeks

**Fig. S11.**
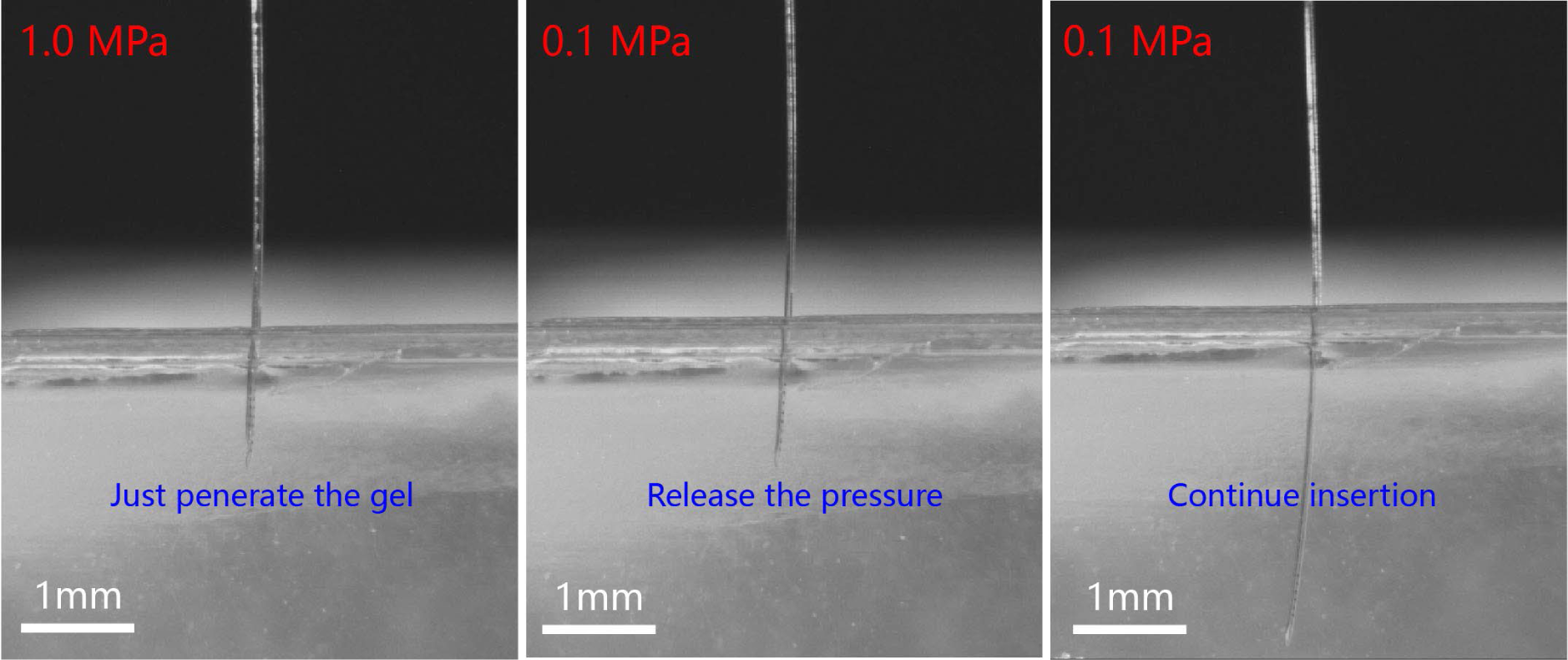
Insert the neurotentacles into the brain-like gel with varied pressure

**Table S1.** Uncalibrated cross-sectional parameters of a neurotentacle.

| **Pressure**<br>**(MPa)** | **Width ($\mu\text{m}$)** | **Thickness ($\mu\text{m}$)** | **Area ($\mu\text{m}^2$)** | **Circumference ($\mu\text{m}$)** |
| --- | --- | --- | --- | --- |
| **0.0** | 182.4 | 9.9 | 1404.0 | 370.0 |
| **0.1** | 192.2 | 17.6 | 1961.2 | 394.6 |
| **0.2** | 162 | 45.6 | 3391.4 | 372.8 |
| **0.5** | 154.9 | 54.4 | 4053.2 | 375.7 |
| **1.0** | 147.9 | 58.6 | 4014.8 | 372.6 |

**Movie S1** Implantation process of the neurotentacle

**Movie S2** Critical implantation pressure testing of neurotentacles on the brain-like gel

**Movie S3** Inserting neurotentacles with variable stiffness in the brain-like gel

